# ORF1ab Codon Frequency Model Predicts Host-Pathogen Relationship in Orthocoronavirinae

**DOI:** 10.1101/2025.01.17.633649

**Authors:** Phillip E. Davis, Joseph A. Russell

## Abstract

Predicting phenotypic properties of a virus directly from its sequence data is an attractive goal for viral epidemiology. Here, we focus narrowly on the Orthocoronavirinae clade and demonstrate models that are powerfully predictive for a human-pathogen phenotype with >86% average precision and >99% average recall on the withheld test set groups, using only Orf1ab codon frequencies. We show alternative examples for other viral coding sequences and feature representations that do not perform well and discuss what distinguishes the models that are performant. These models point to a small subset of features, specifically 5 codons, that are critical to the success of the models. We discuss and contextualize how this observation may fit within a larger model for the role of translation in virus-host agreement.

## Introduction

There are several examples of modeling efforts attempting to assess either the host range or zoonotic potential of RNA viruses broadly, and Coronaviruses specifically in the wake of COVID-19 [1-5].

There are many commonalities between these approaches. Naturally, due to its role in receptor binding, Spike protein is often the focus of these efforts when analyzing properties such as nucleotide or amino acid composition. Limitations to focusing on Spike protein exclusively have been described previously [3,6]. Other approaches look at compositional biases in the complete viral genome or proteome.

Compositional representations are also usually drawn from a short list of possibilities such as dinucleotide composition, or a measure of codon bias such as Relative Synonymous Codon Usage (RSCU). The motivations for these representations comes from observed compositional biases across a variety of viral families [7]. Especially in the case of codon composition, many computational and experimental results point to the importance of host-virus agreement in the translational environment. For example, virus codon compositions are predictive of their tissue tropism [8], and strategies have been discovered in which both the host and viruses manipulate cellular tRNA abundances to either restrict or promote virus replication [8-12]. However, this phenomenon has not yet been leveraged for a pathogen-class predictive model to our knowledge. Here, we present our results applying codon frequencies as the feature space for a model capable of distinguishing human-pathogen Orthocoronavirinae examples from non-human-pathogens.

## Results

We report that models fit on Orf1ab with a simple codon frequency feature representation predict accurately (average 88.6%) on sequences from withheld species groups. Model recall was especially high (average 99.7%), with false negatives being nearly entirely absent (**Figure 1**). The exception to this is when Betacoronavirus 1 is the negative test case, where all models had difficulty predicting the negative class label. Our results point to two interesting findings — 1) using only codon frequency as the feature representation results in a boost in performance over RSCU across both recall and precision, and 2) depending on the feature representation, information content about host-pathogen potential can vary dramatically across the genome. As far as we know, these are the only modeling results reported that have relied solely on the Orf1ab coding region.

**Figure 1.**
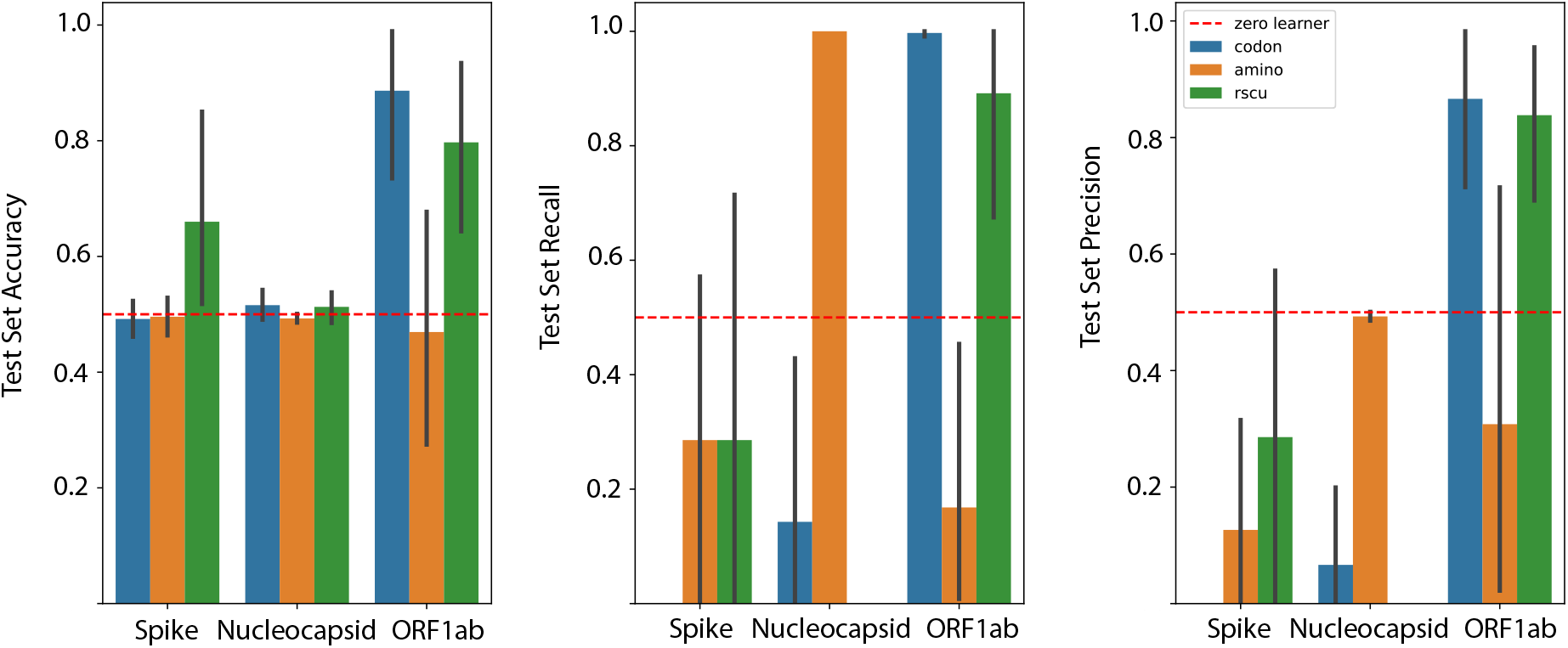
Average performance metrics across each of the seven test set splits for each combination of viral CDS and feature representation. Codon frequency model is top performer, with boosts in average performance across each metric over RSCU.

Since L1-regularization provides feature selection, this allows for interpretation of the model results. Inspecting the codon frequency models shows that models selected by our cross-validation routine are consistently sparse, using between 7 and 13 of the possible 64 codons. Of these, 5 codons have non-zero coefficients in every model fit, no matter what test set group was withheld. These are Thr^ACG^, Ser^TCG^, Trp^TGG^, Gln^CAA^, and Ala^GCT^ (**Table 1**).

**Table 1.** Predictors from each of the codon frequency models across the seven test set splits with non-zero coefficients. Positive coefficients are highlighted in green while negative coefficients are highlighted in red. We identified 5 codon predictors which appear with non-zero coefficients in every model (bold text / red boxes), no matter the withheld test set data.

| Codon | 229E<br>and MHV | HKU1<br>and PEDV | MERS and<br>AlphaCOV1 | NL63 and<br>BetaCOV1 | OC43 and<br>Avian COV | SARS-CoV-2<br>and SADS | SARS-CoV-1<br>and HKU15 |
| --- | --- | --- | --- | --- | --- | --- | --- |
| **ACG** | **-0.4886** | **-0.5521** | **-0.6272** | **-0.3837** | **-0.6223** | **-1.1190** | **-0.4819** |
| **TCG** | **-0.2666** | **-0.3789** | **-0.4251** | **-0.3427** | **-0.5375** | **-0.0727** | **-0.5009** |
| ACT | 0.1713 | 0.1856 | 0.1824 | 0.0000 | 0.4700 | 0.4240 | 0.3808 |
| TTA | 0.1761 | 0.0000 | 0.1337 | 0.0108 | 0.0000 | 0.0000 | 0.1703 |
| **TGG** | **0.2238** | **0.5095** | **0.6464** | **0.4319** | **0.8253** | **0.8400** | **0.7919** |
| **CAA** | **0.2944** | **0.6909** | **0.6361** | **0.5715** | **0.7150** | **0.5419** | **0.7591** |
| **GCT** | **0.4536** | **0.5278** | **0.3378** | **0.7633** | **0.3290** | **0.4261** | **0.2325** |
| CCG | 0.0000 | -0.1530 | -0.1406 | -0.0857 | 0.0000 | 0.0000 | 0.0000 |
| GCA | 0.0000 | -0.0373 | 0.0000 | 0.0000 | 0.0000 | 0.0000 | -0.1479 |
| GGG | 0.0000 | 0.0000 | -0.1012 | 0.0000 | -0.0246 | 0.0000 | 0.0000 |
| TAA | 0.0000 | 0.0000 | 0.0007 | 0.0000 | 0.0000 | 0.0000 | 0.0000 |
| GAT | 0.0000 | 0.0000 | 0.0656 | 0.0000 | 0.0000 | 0.0000 | 0.0000 |
| ATT | 0.0000 | 0.0000 | 0.1794 | 0.0000 | 0.0000 | 0.1464 | 0.1274 |
| AGA | 0.0000 | 0.0000 | 0.2377 | 0.0000 | 0.0000 | 0.0000 | 0.0000 |
| AGC | 0.0000 | 0.0000 | 0.0000 | 0.0000 | -0.1565 | 0.0000 | 0.0000 |
| CTC | 0.0000 | 0.0000 | 0.0000 | 0.0000 | 0.0672 | 0.0000 | 0.0000 |
| AGT | 0.0000 | 0.0000 | 0.0000 | 0.0000 | 0.0000 | 0.0000 | 0.0520 |
| TCT | 0.0000 | 0.0000 | 0.0000 | 0.0000 | 0.0000 | 0.0000 | 0.0890 |

The appearance of the tryptophan codon as a top predictor in this model (*which is not a feature of RSCU models as there is only a single codon for tryptophan*) indicates a source of the performance gain. In contrast, while tryptophan amino acid frequency is available to the amino acid models, these models did not perform well on withheld data. This performance advantage is only realized when modeling the interaction between codon frequency features. To illustrate the importance of the tryptophan codon feature, **Table 2** shows the non-zero-coefficient predictors in the codon frequency and RSCU models. Several predictors overlap between the two (indicated in bold), with the strongest predictor in the codon frequency model being Trp^TGG^.

**Table 2.** Predictors and coefficients for model fit without SARS-CoV-2 and SADS (Test Set Pair #6) in both the codon frequency and RSCU representation.

| Codon Frequency |  | RSCU |  |
| --- | --- | --- | --- |
| **ACG** | -1.1190 | **ACG** | -0.4840 |
| **TCG** | -0.0727 | **CCG** | -0.2987 |
| **ATT** | 0.1464 | **GCA** | -0.2788 |
| **ACT** | 0.4240 | **TCG** | -0.2597 |
| **GCT** | 0.4261 | **CTA** | -0.1558 |
| **CAA** | 0.5419 | **CAG** | -0.0246 |
| **TGG** | 0.8400 | **CAA** | 0.1565 |

## Methods

### Data Collection

Sequence data for this study was gathered from the NCBI Virus database for sequences submitted before December 2019. For SARS-CoV-2, sequences were downloaded and subsampled to include 10 records from each variant of concern (VOCs) as of August 2021: Alpha, Beta, Gamma, and Delta. Viral coding sequences (CDS) were categorized by gene, resulting in datasets for Orf1ab (N=2270), Spike (N=2198), and Nucleocapsid (N=2201).

### Pathogen Classification

Sequences were labeled as **human-pathogenic** if they were associated with known disease in humans. This classification included seven positive human-pathogen groups: Human CoV 229E, Human CoV HKU1, MERS-CoV, Human CoV NL63, Human CoV OC43, SARS-CoV-1, and SARS-CoV-2. Non-human-pathogenic representatives included various Alphacoronaviruses, Betacoronaviruses, Gammacoronaviruses, and Deltacoronaviruses. These sequences were labeled as non-human-pathogens unless they caused disease in humans. For example, SARS-CoV-1 isolates from civets and MERS isolates from camels were classified as human pathogens, while all other isolates remained non-human-pathogens.

### Cross-Validation and Test Set Design

The primary goal was to evaluate the model’s ability to generalize to novel species within the Orthocoronavirinae clade. To achieve this, a group-based cross-validation strategy was employed. Each sequence was assigned a group label based on its species-level taxonomy ID and pathogen class (human or non-human). For instance; Human CoV OC43 was labeled as “1_694003” (human pathogen representative of Betacoronavirus 1), while Bovine CoV and Porcine Hemagglutinating Encephalomyelitis Virus (PHEV) were labeled as “0_694003” (non-human-pathogen representatives of Betacoronavirus 1). Seven test set splits were pre-defined, with each split excluding one human-pathogen species as a positive test set and one non-human-pathogen species as a negative test set. This ensured that the test set always represented novel species not seen during training. The test set pairs were as shown in **Table 3**.

**Table 3.**
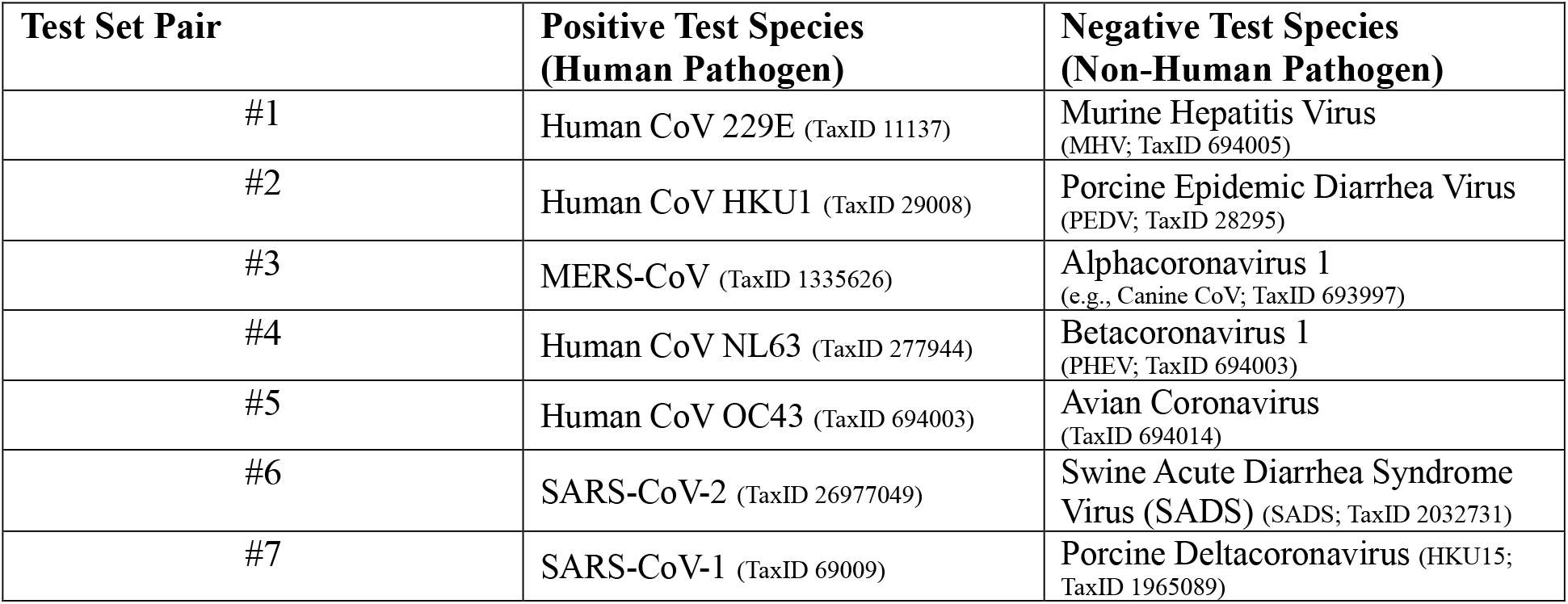
Human-pathogen and non-human-pathogen pairs selected for test set splits.

### Data Balancing and Supersampling

For each test set split, the remaining sequences were divided into training and validation sets. Training data were supersampled to ensure balance between human and non-human-pathogen classes. A two-tiered randomization process first determined whether to sample from the positive or negative class, then uniformly sampled sequences from the corresponding groups. Training data were supersampled to 4000 records, while test sets were subsampled to 200 records.

### Feature Representations

Three feature extraction methods were tested:

1. ***Amino acid frequency***: Counts of each amino acid in the CDS.
2. ***Relative Synonymous Codon Usage (RSCU)***: Codon counts normalized by the number of synonymous codons for each amino acid.
3. ***Codon frequency***: Raw counts of each codon, independent of synonymous grouping.

### Model Training and Selection

All models were trained using L1-regularized logistic regression. To prevent data leakage, preprocessing steps (e.g., standard scaling) were applied within a scikit-learn pipeline. Cross-validation was performed using a leave-two-groups-out strategy, where one positive and one negative group were withheld for validation. Hyperparameters, including the regularization term and model tolerance, were optimized using randomized cross-validation with 1000 parameter combinations sampled.

### Model Evaluation

Model performance was assessed using metrics such as accuracy, precision, recall, and the negative Brier score. We averaged the performance statistics for accuracy, recall and precision across each of the seven test and training set splits for each combination of viral CDS and feature representation.

Persistent model objects for the selected model, a cross-validation report, a list of training set and test set misclassified records, a summary of model parameters, and a table of the non-zero coefficient predictor variables and their coefficients were saved with each model fit for each test set split and are available at the GitHub repository along with the code used to produce the models (https://github.com/mriglobal/codon_amino_cov/tree/main).

## Discussion

These results provide new insights into assumptions underlying viral genotype-to-phenotype modeling. While the specific biological phenomena driving the improved performance of codon frequency models and the focus on Orf1ab remain uncertain, the interpretation of these models offers potentially valuable context. First, as previously noted, the inclusion of tryptophan codon frequency emerged as a key factor. In logistic regression models, the coefficients for each predictor reflect the increase or decrease in the log likelihood of the positive class for each unit of the predictor variable. Predictors with positive coefficients are enriched in the positive class, whereas those with negative coefficients are depleted.

Notably, ThrACG and SerTCG consistently exhibited negative coefficients across all codon frequency models. These codons are the rarest for their respective amino acids and are among the rarest codons in the human genome, which may explain their significance in the models. Numerous theoretical factors contribute to codon usage bias in viruses, including CpG avoidance for Zinc finger antiviral protein binding [13], RNA secondary structure requirements [14], and alignment with host translational preferences [15]. While these models cannot directly elucidate underlying mechanisms, they provide hypotheses for the observed patterns. Relative synonymous codon usage (RSCU) is commonly employed to model translation-influenced systems, such as gene expression [16]. However, in a tRNA supply-and-demand framework, RSCU is insensitive to codon compositional features, such as tryptophan codon frequency, that may influence translation efficiency. Evidence suggests that viruses can manipulate host tRNA pools to enhance replication. For example, Flaviviruses counteract Schlafen-family viral restriction genes [9], HIV alters tRNA abundances to improve translation efficiency [17], and Influenza A and Vaccinia viruses modulate translationally active tRNA pools at the polysome [18]. Whether Schlafen-family genes are activated during coronavirus infections or whether TRMT-1 cleavage by the main protease [19] affects cellular tRNA abundances remains unknown. However, data indicate that both tRNA levels and 5-methoxycarbonylmethyl-2-thiouridine modifications of certain position-34-U tRNA isoacceptors are enriched in SARS-CoV-2-infected cells [20], including the GlnCAA codon identified in our models as a significant predictor.

Compositional representations of viral sequences likely reflect aggregated effects from these biological factors. However, assuming uniform selective pressures across viral families introduces a strong limitation. While there may be general compositional similarities among human-infecting viruses, increased predictive power in genotype-to-phenotype models can be achieved through enhanced experimentation and feature engineering. The models presented here demonstrate substantial information content regarding human-pathogenic potential in specific genomic contexts. However, they remain incomplete, as many known barriers to zoonosis in coronaviruses are not addressed. These barriers include, for example, cross-reactivity to endogenous human coronaviruses in the case of AlphaCoV1 [21] and 229E-related Camel alphacoronaviruses [22], as well as host-receptor binding compatibility. Nevertheless, the findings suggest that ensemble models incorporating these approaches could improve predictive power in coronaviruses. More broadly, general genotype-to-phenotype modeling efforts in viruses could benefit from similar strategies.

